# Xist Intron 1 Repression by TALE Transcriptional Factor Improves Somatic Cell Reprogamming in Mice

**DOI:** 10.1101/259234

**Authors:** Jindun Zhang, Xuefei Gao, Jian Yang, Xiaoying Fan, Wei Wang, Yanfeng Liang, Lihong Fan, Hongmei Han, Xiaorong Xu, Fuchou Tang, Siqin Bao, Pentao Liu, Xihe Li

**Affiliations:** Research Center for Animal Genetic Resources of Mongolian Plateau, Inner Mongolia University, Hohhot, 010021, China; Wellcome Trust Sanger Institute, Cambridge CB10 1HH, United Kingdom; Biodynamic Optical Imaging Center (BIOPIC), Peking University, Beijing, 100871, China; College of Life Sciences, Peking University,Beijing, 100871, China; Inner Mongolia Saikexing Institute of Breeding and Reproductive Biotechnology in Domestic Animal, Hohhot, 011517, China

**Author notes:** Correspondence should be addressed to Xihe Li, Ph.D., or Pentao Liu, Ph.D.

**Keywords:** *Xist* intron 1, TALE dTFs, MEF reprogramming, iPSC, SCNT

## Abstract

*Xist* is the master regulator of X chromosome inactivation (XCI). In order to further understand the *Xist* locus in reprogramming of somatic cells to induced pluripotent stem cells (iPSCs) and in somatic cell nuclear transfer (SCNT), we tested transcription-factor-like effectors (TALE)-based designer transcriptional factors (dTFs), which were specific to numerous regions at the *Xist* locus. We report that the selected dTF repressor 6 (R6) binding the intron 1 of *Xist*, which did not affect *Xist* expression in mouse embryonic fibroblasts (MEFs), substantially improved the iPSC generation and the SCNT preimplantation embryo development. Conversely, the dTF activator targeting the same genomic region of R6 decreased iPSC formation, and blocked SCNT-embryo development. These results thus uncover the critical requirement for the *Xist* locus in epigenetic resetting, which is not directly related to *Xist* transcription. This may provide a unique route to improving the reprogramming.

## Introduction

In the mouse, the two X chromosomes are active in the epiblasts of the blastocyst as well as in pluripotent stem cells. In subsequent embryo development or upon differentiation of pluripotent stem cells, one X chromosome becomes inactivated to balance the expression of X-linked genes between male and female cells [1]. The X chromosome inactivation (XCI) process is triggered by *Xist* transcript coating the X chromosome that becomes silent [2–4].

X chromosome inactivation and reactivation are a dynamically-regulated developmental process which is associated with loss or reacquisition of pluripotency in embryonic stem cell (ESC) differentiation, or in reprogramming of somatic cells to iPSCs or in SCNT. Previous SCNT studies indicate that the inactivated X chromosome is frequently improperly reactivated as it exhibits the epigenetic memory of the parental X chromosome [5–8]. It has also been reported that, the X-linked genes are down-regulated in many cloned embryos, and that the *Xist* expression levels are significantly higher in cloned embryos (both male and female) than in control IVF embryos [9]. The functional significance of the *Xist* in SCNT has been demonstrated in recent studies where deletion of the *Xist* gene on the active X chromosome (Xa) or Xist-siRNA results in relatively normal global gene expression in SCNT-derived preimplantation embryos and yields an eight- to nine-fold increase in cloning efficiency in terms of live birth rates [9–10].

Reprogramming of female somatic cells to iPSCs by the four transcription factors *Oct4, Sox2, Klf4*, and *c-Myc* (OSKM) also leads to X reactivation [11]. The transition from pre-iPSC in the late phases of reprogramming to full pluripotent iPSCs requires repression of *Xist* expression [12–13], whereas knockdown of *Xist* improves the efficiency in late stages of female MEF reprogramming [14]. When histone variants TH2A/TH2B that are usually found in oocytes and early embryos are ectopically expressed together with OSKM, the reprogramming efficiency is 2.4-3.6 fold higher from *Xist* mutant MEFs than that from wild type (WT) cells [15].

In ESCs, pluripotency factors OCT4, SOX2 and NANOG bind a genomic region in intron 1 of the *Xist* locus, which was thought to repress *Xist* expression [16]. Additionally, other pluripotency regulators such as TCF3 and PRDM14, and the early developmental regulators such as CDX2, also bind the same region, which is thus considered as an enhancer [17–20]. Surprisingly, despite of the binding of numerous transcription factors, this enhancer does not appear to function as an active enhancer in undifferentiated ESCs [21], and is dispensable for X chromosome inactivation, reactivation or for critical control of *Xist* expression [22]. Instead, it is proposed that it functions in ESC differentiation [22].

Transcriptional-activator-like effectors (TALEs) based designer transcriptional activators (A-dTFs) and repressors (R-dTFs), which fuse TALE proteins for specific genomic DNA sequences with an activation or a repression domain, enable epigenetic modifications of a chosen genomic locus and regulate its expression [23]. dTFs have been tested in MEF reprogramming, directed ESC differentiation and transdifferentiation [24–27].

In this study, we made and expressed dTFs that specifically bound several genomic regions at the *Xist* locus, and subsequently examined their functions in reprogramming somatic cells to iPSCs and in SCNT. Unexpectedly, the dTF repressor binding the intron 1 enhancer region substantially improved iPSC production and SCNT preimplantation embryonic development correlated with much fewer abnormally expressed genes frequently associated with SCNT, even though it did not affect *Xist* expression. In stark contrast, the dTF activator targeting the same enhancer region drastically decreased both iPSC generation and SCNT efficiencies and induced ESC differentiation. Our results thus uncover a previously unrecognized role of the *Xist* locus in epigenetic reprogramming.

## Results

### Xist expression in the presence of dTF repressors

We designed and made several TALE proteins that were specific to DNA sequences of the *Xist* locus utilizing a TALE repeat library [28]. These genomic sequences were from the promoter region and the intron 1 enhancer region (Figure 1A and Table EV1). In the dTF designs, the TALE proteins were fused with the repression domain KRAB [25], which were linked to mCherry by the T2A peptide for convenient tracking of their expression in cells (Figure EV1A). Expression of the dTF repressors was controlled by the Tet/On system where doxcycline (Dox) induces gene expression. The expression cassettes of the dTF repressors were delivered to cells by *piggyBac* (PB) transposition [29].

**Figure 1.**
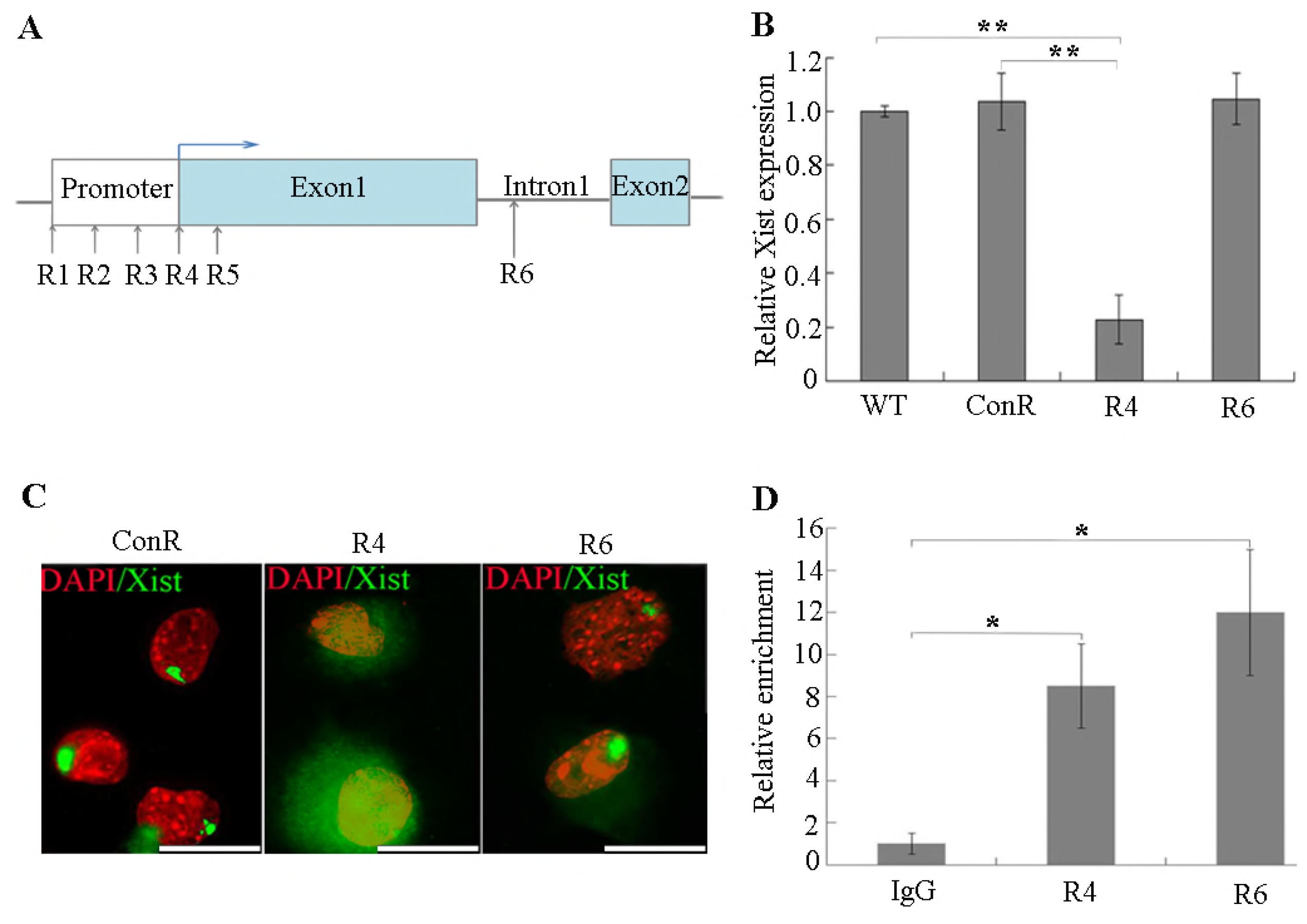
Schematic for dTFs targeting *Xist* locus and characterization of *Xist* regulation. (A) The binding sites of TALE dTFs at the *Xist* locus. (B) qRT-PCR analysis of expression of *Xist* in female MEFs on D5. (C) RNA-FISH analysis of *Xist* (green) in female MEFs on D5. Scale bars=10 μ m. (D) Validation of TALEs, in female cells, binding to the *Xist* locus in the ChIP assay using an antibody against HA tag followed by qPCR to amplify the corresponding genomic DNAs. Primer pair 3 from the *Xist* intron 1 was used, and IgG was used as the control. Results are representative of three independent lines and are mean ± SD. n = 3. **p < 0.01. *p <0.05.

We next investigated the effects of expressing dTF repressors on *Xist* expression. Female MEFs were transfected with the PB transposons carrying the dTFs and with a PB transposase expression plasmid, to facilitate integration of the dTF expression cassettes into the genome for stable expression. Dox was added to induce dTF repressor expression, and the transfected cells were collected five days after transfection for analysis. Quantitative reverse transcription-PCR (qRT-PCR) analysis revealed that among those dTF repressors that bind the promoter/exon 1 region (R1, R2, R3, R4, R5), R4 caused substantial decrease of *Xist* compared to that in either wild type female MEFs or those expressing a dTF repressor specific to an unrelated genomic region (ConR) (Figure 1B and Table EV1; *P*<0.01). In contrast, R6 that binds the intron 1 enhancer region did not cause any noticeable changes in *Xist* expression (Figure 1B), in line with the previous study where deletion of this region did not affect *Xist* expression [22]. We confirmed *Xist* expression at the single-cell level by fluorescence in situ hybridization (FISH) where a strand-specific RNA probe to *Xist* was used to detect and to localize *Xist* transcript. In female MEFs expressing R4, approximately 60% of them displayed virtually no *Xist* RNA cloud or pinpoint signal, whereas the rest cells had a faint *Xist* signal (Figure EV1B). In line with the qRT-PCR results, cells expressing R6 or the ConR showed comparable *Xist* FISH signals as in the wild type female cells (Figure 1C and Figure EV1C). The failure of R6 to regulate *Xist* expression might be caused by the low binding affinity of R6 to its target sequence in the intron 1 enhancer region. To exclude this possibility, we replaced the KRAB domain in the dTF repressors with the 2 × HA hemagglutinin (HA) tag (Figure EV1A), and performed ChIP-qPCR analysis using female cells expressing the HA-tagged TALEs. The tagged TALE proteins of R6 and R4 showed comparable binding enrichments for their respective target sequences (Figure 1D and Table EV2; *P*<0.05). Thus, the difference of regulating *Xist* expression by R4 and R6 is not caused by their distinct binding affinities to the targets.

### dTF repressors improve reprogramming of MEFs to iPSCs

In order to examine the effects of dTF repressors on reprogramming of somatic cells to iPSCs, we co-expressed by Dox induction OSKM along with R4 or R6 in female or male *Oct4*-GFP reporter MEFs [30]. All expression cassettes were delivered by the PB transposition [31]. Two weeks after transfection, Dox was removed and the medium was switched to N2B27/2i/LIF, which allows selection for naïve iPSC colonies (Figure EV2A). One week later, ESC-like colonies started to emerge, and eventually Dox independent iPSC colonies formed (Figure 2A). Co-expressing the ConR with the four factors produced similar numbers of iPSC colonies as in the control where no dTF was expressed (Figure 2B). In contrast, co-expression of R4 or R6 with the four factors gave rise to an approximately 2.5- and 3.5-fold of iPSC colonies, respectively, compared to the control (Figure 2B; *P*<0.05). Interestingly, we also obtained substantially more iPSC colonies when R4 or R6 was co-expressed with the four factors in male MEFs (Figure 2B; *P*<0.05). Importantly, R6-coexpression always produced more iPSC colonies than R4 co-expression, indicating that R6 facilitates epigenetic changes independent of *Xist* expression. To confirm pluripotency of the iPSCs produced by expressing dTF repressors, several iPSC lines were characterized by qRT-PCR and differentiation assays *in vitro*, which showed that the iPSCs expressed appropriate levels of pluripotent genes when compared to the control (Figure 2C), and could differentiate to cells of the three germ layers (Figure EV2B). Finally, chimeric mice were derived from these iPSCs confirming their pluripotency in the *in vivo* development (Figure 2D).

**Figure 2.**
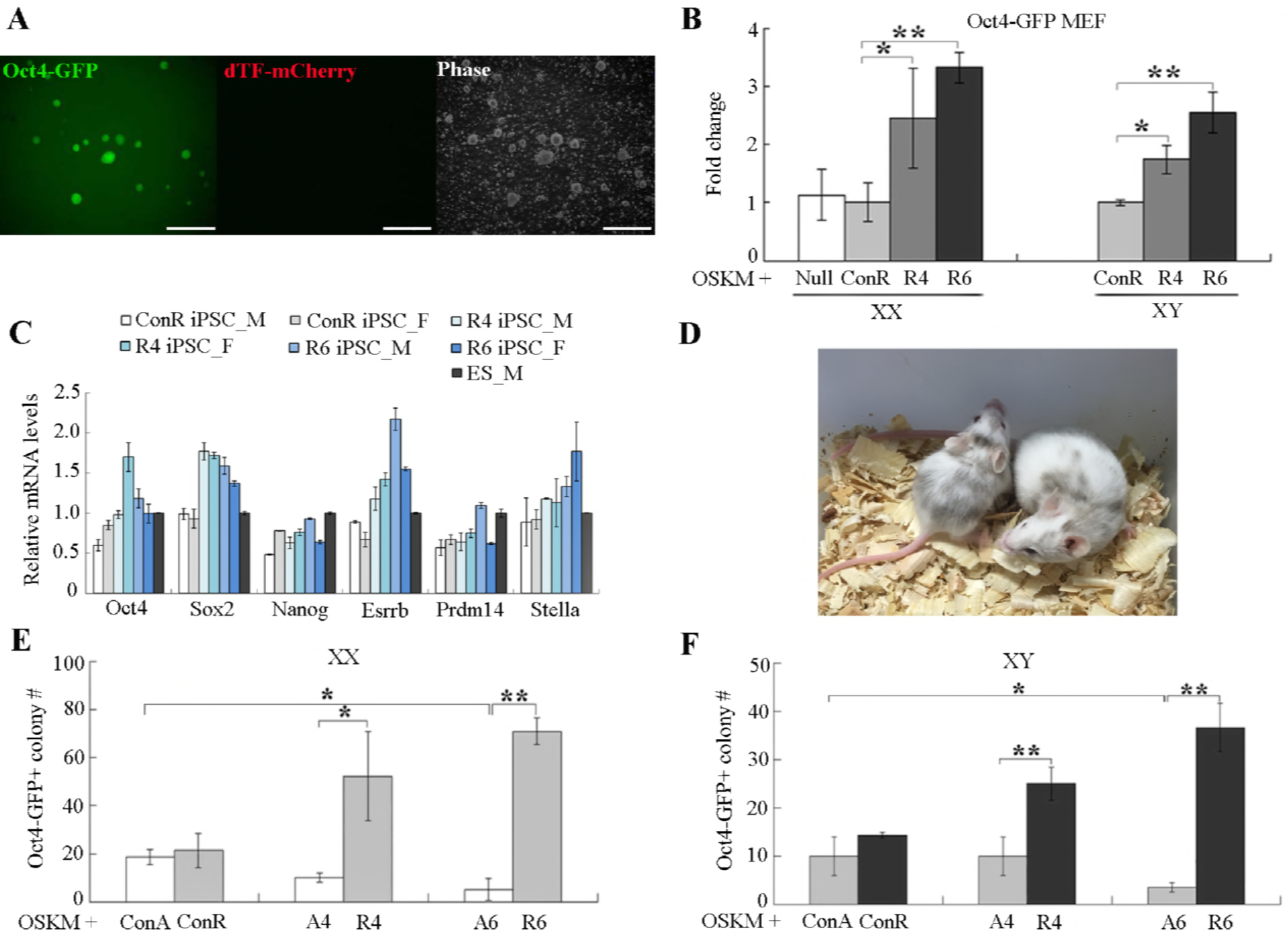
Effects of *Xist* dTFs on iPSC induction. (A) The morphology of Dox-independent iPSCs. (B) The rate of iPSC formation from Oct4-MEFs induced by 4F+R4 and 4F+R6, respectively, compared to 4F+ConR. (C) qRT-PCR analysis for selected ESC markers in established iPSCs. Data are shown relative to GAPDH expression level in male ESCs. (D) Contribution of 4F*+Xist* TALE dTF derived iPSCs to the chimeras. (E and F) The number of female and male Oct4-GFP iPSC colonies produced by 4F+dTFs. Results are representative of three independent lines and are mean ± SD. n = 3. **p <0.01. *p <0.05

We asked whether dTF activators of the *Xist* locus might have an opposite effect on MEF reprogramming. To make dTF activators, we replace the KRAB domain in R4 and R6 with the VP64 domain to make A4 and A6, respectively (Figure EV1A). The ConA control activator was constructed from ConR. We co-expressed OSKM with either R4, R6 or A4, A6 in *Oct4*-GFP MEFs for iPSC generation (Figure EV2A). In female *Oct4*-GFP iPSC induction, co-expression of A4 produced drastically (2-folds) fewer iPSC colonies in comparison to ConA co-expression (Figure 2E), whereas A6 expression reduced iPSC colonies by 4 folds compared to ConA (Figure 2E; P<0.05). Similarly, expressing dTF activators A6 also compromised reprogramming of male MEFs to iPSCs (Figure 2F; P<0.05). The number of AP staining-positive colonies was consistent with the *Oct4*-GFP results (Figure EV2C).

The effects of dTFs on MEF reprogramming were confirmed by reprogramming *Rex1*-GFP reporter MEFs [32] to iPSCs (Figure EV2D and Figure EV2E; *P*<0.01), especially when co-expressing R6 or A6. The opposing effects of expressing dTF repressors or activators for the *Xist* intron 1 enhancer demonstrate the importance of this region in epigenetic reprogramming, which is likely unrelated to *Xist* transcript levels from R6/A6 data.

### R6 improves development of SCNT preimplantation embryos

SCNT is frequently associated with dysregulation of a large number of genes but only a small number of these genes are common across different cloned embryos [9]. Surprisingly, 54% of the commonly down-regulated genes are X-linked genes [9]. Moreover, *Xist* is found to be frequently ectopically expressed which is associated with aberrantly X inactivation in both male and female cloned embryos [7–9], Deletion of the *Xist* exon 1 or knocking-down *Xist* substantially increases SCNT efficiency in terms of live born animals [9, 10]. Since R4 and R6 affected MEF reprogramming to iPSCs, we investigated whether they might also improve SCNT.

In order to enrich and select for MEFs that expressed the dTF repressors as SCNT cloning donors, we co-transfected MEFs with a plasmid carrying the Puro-IRES-GFP cassette along with Dox-inducible dTF repressor (mCherry) cloned in PB transposons. The transfected MEFs were briefly selected with puromycin, and the GFP^+^/mCherry^+^ MEFs (GFP/mCherry) were used as donor cells in SCNT (Figure EV3A and Figure 3A). The reconstructed embryos were cultured in KSOM medium for 96 hours to allow preimplantation development (Figure EV3A and Figure 3B). When wild type female donor MEFs were used, 5.8% cloned 2-cell embryos developed to the blastocyst stage (Figure EV3B and Table EV3). When ConR was expressed in female MEFs, this percentage was at about 2.7% (Figure EV3B and Table EV3). If R6 was expressed in the female donor cells, the developmental rate of the cloned 2-cell embryos to blastocysts was 9.5% (Figure EV3B and Table EV3; *P*<0.01). Also, R6 improved the male SCNT-blastocyst development from 2-cell embryos (Figure EV3C and Table EV3; P<0.01). Remarkably, R6 further considerably enhanced the development from morulae to blastocysts in both female and male embryos (Figure 3C, 3D, and Table EV3; *P*<0.01): for embryos from wild type female MEFs or the ones expressing ConR, the development rates were 17.6% and 17.4%, respectively, and R6 expression increased the rate to 53.3% (Figure 3C; *P*<0.01). In the case of male MEF donors, expressing R6 increased the morula/blastocyst rate from 50.4% to 91.5% (Figure 3D; *P*<0.01). These data indicate that R6, although it did not affect *Xist* expression, substantially improved both reprogramming of somatic cells to iPSCs and SCNT. By contrary, SCNT embryos from those MEFs expressing A6, the dTF activator for the intron 1 enhancer, almost completely failed in preimplantation development (Table EV3). On the other hand, expressing R4, which down-regulated *Xist* expression, did not significantly alter the developmental potential of the cloned embryos (Table EV3), which is consistent with the reports that neither *Xist* deletion nor *Xist* knockdown improved preimplantation development of SCNT embryos [9, 10].

**Figure 3.**
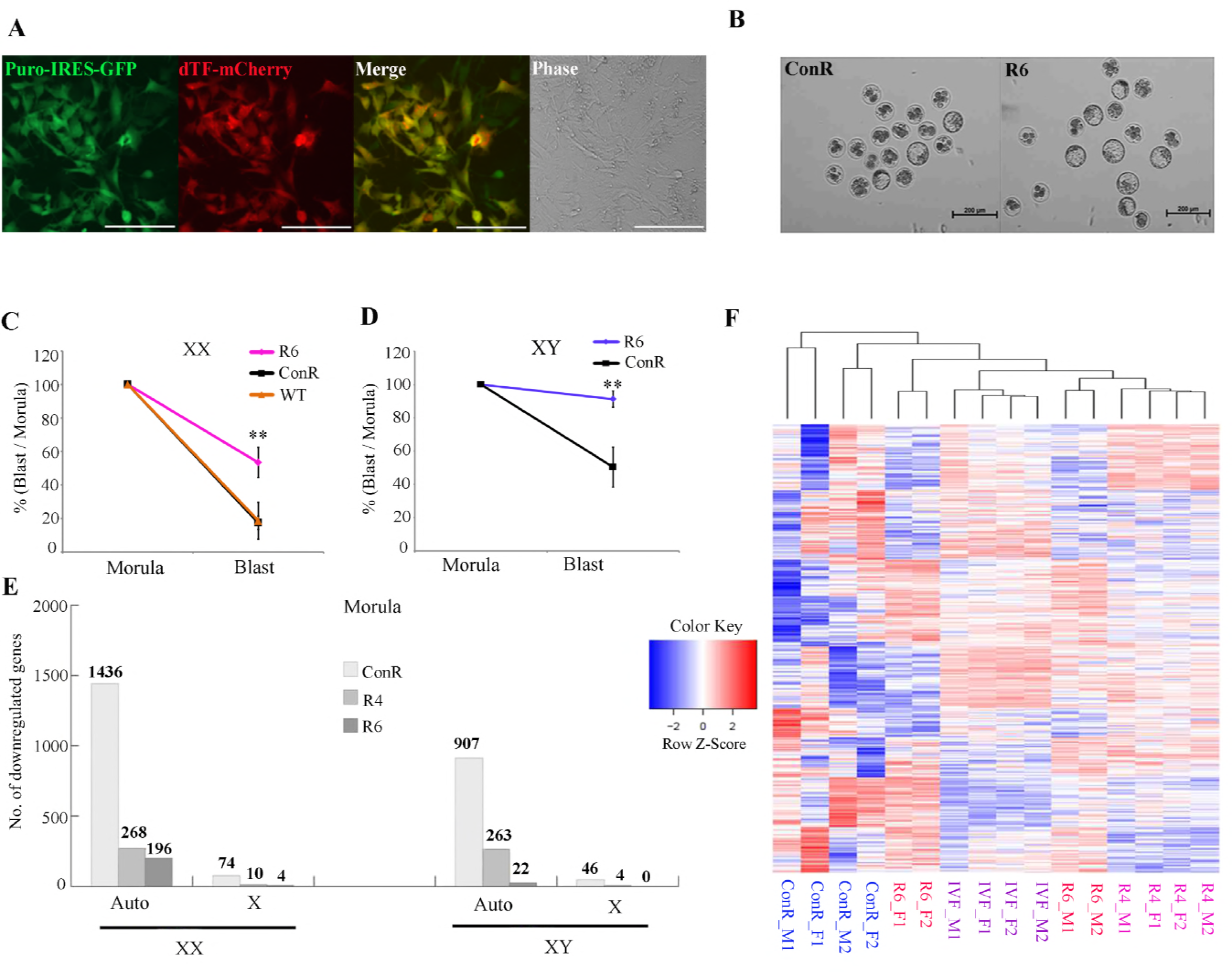
XistdTF repressors improve the developmental potential of SCNT embryos. (A) Morphology of transgenic donor MEFs expressing both Puro-IRES-GFP and *Xist* dTF-mCherry. Scale bar=200 μ m. (B) Representative images of female SCNT embryos after 96 h of *in vitro* culture. Scale bars =200 μ m. (C and D) The percentage of female and male cloned blastocysts, respectively, that developed from morula stage embryos *in vitro*. (E) Bar chart illustrating the reduction in the number of differentially expressed genes (FC > 10) between IVF and SCNT morulae after *Xist* dTF repressor transfection. (F) Hierarchical clustering of samples used in the study. **p <0.01.

We subsequently investigated whether R6 altered gene expression in SCNT preimplantation embryos, which might account in part for its ability to improve SCNT. To address this possibility, individual SCNT morulae from MEFs expressing dTF repressors (R6 or ConR) were harvested for RNA sequencing (RNA-seq). The gene expression profiles of the SCNT embryos were compared to that of morulae from *in vitro* fertilization (IVF; FPKM > 5). Remarkably, compared to morulae from female MEFs expressing ConR, R6 expression led to drastic decrease of the commonly down-regulated X-linked genes from 74 to 4 (Figure 3E, FC > 10). In embryos from male MEFs, SCNT caused dysregulation of a large number of X-linked genes, but most of them were embryo-specific. Remarkably, no commonly down-regulated X-linked gene was found when R6 was expressed (Figure 3E, FC > 10). Furthermore, the number of autosomal genes commonly down-regulated in the morulae from female or male MEFs expressing R6 declined by 86.3% (1436 to 196) and 97.6% (907 to 22), respectively, compared to that in ConR morulae (Figure 3E, FC > 10). Hierarchical clustering transcriptome analysis of the RNA-seq data revealed that the transcriptomes of SCNT morulae from male MEFs expressing R6 were more closed clustered with those of IVF embryos than ones from MEFs expressing ConR (Figure 3F and Figure EV3D), and this effect was more significantly than female embryos (Figure 3F and Figure EV3E). Therefore, R6 expression improves the SCNT preimplantation embryo development possibly by regulating expression of a number of X-linked and autosomal genes.

### A6 promotes ESC differentiation

A6, as is shown above, prevented the MEF reprogramming, indicating that it could cause ESC differentiation. We first expressed A6 or ConA in male *Oct4*-GFP ESCs. Expression of A6 appeared to reduce *Oct4* since the mCherry^+^ cells (expressing A6) became GFP^dim^ or GFP^−^ as early as three days after Dox induction (Figure 4A). On the other hand, ConA had no significant effect because the mCherry^+^ ESCs remained GFP^+^ (Figure 4A). We subsequently collected mCherry^+^ ESCs after five days of A6 expression, and quantitated pluripotent gene expression. *Oct4* mRNA levels were decreased substantially in A6 cells, compared to those expressing ConA (Figure 4B; P<0.05). *Prdm14*, which maintains mouse ESCs partly through repression of differentiation and accelerates epigenetic reprogramming of human and mouse somatic cells to iPSCs [19, 33–34], was also reduced in ESCs expressing A6 (Figure 4B; P<0.01). It has been shown that *Prdm14* down-regulates *Dnmt3a* and *Dnmt3b* in ESCs [34]. Consistent with lower *Prdm14* in A6-expressing ESCs, *Dnmt3a* was substantially upregulated (Figure 4B; P<0.01). Concomitantly, lineage marker genes including *Gata6, T* and *Lefty* were also upregulated in ESCs expressing A6 (Figure 4B; P<0.05). Similar to in the male cells, expressing A6 in female ESCs substantially decreased expression of pluripotent genes *Rex1, Oct4* and *Prdm14* (Figure 4C; P<0.05). To further investigate the influence of A6 on ESC differentiation, Oct4-GFP ESCs expressing A6 or ConA were cultured in ESC medium without LIF. After 10 days, A6 expression appeared to accelerate the reduction of fluorescence intensity in *Oct4*-GFP ESCs than ConA expression (Figure 4D). Consequently, the effects of A6 on ESC differentiation demonstrate the importance of *Xist* intron 1 in epigenetic reprogramming, which indirectly confirm the role of R6 in promoting MEF reprogramming.

**Figure 4.**
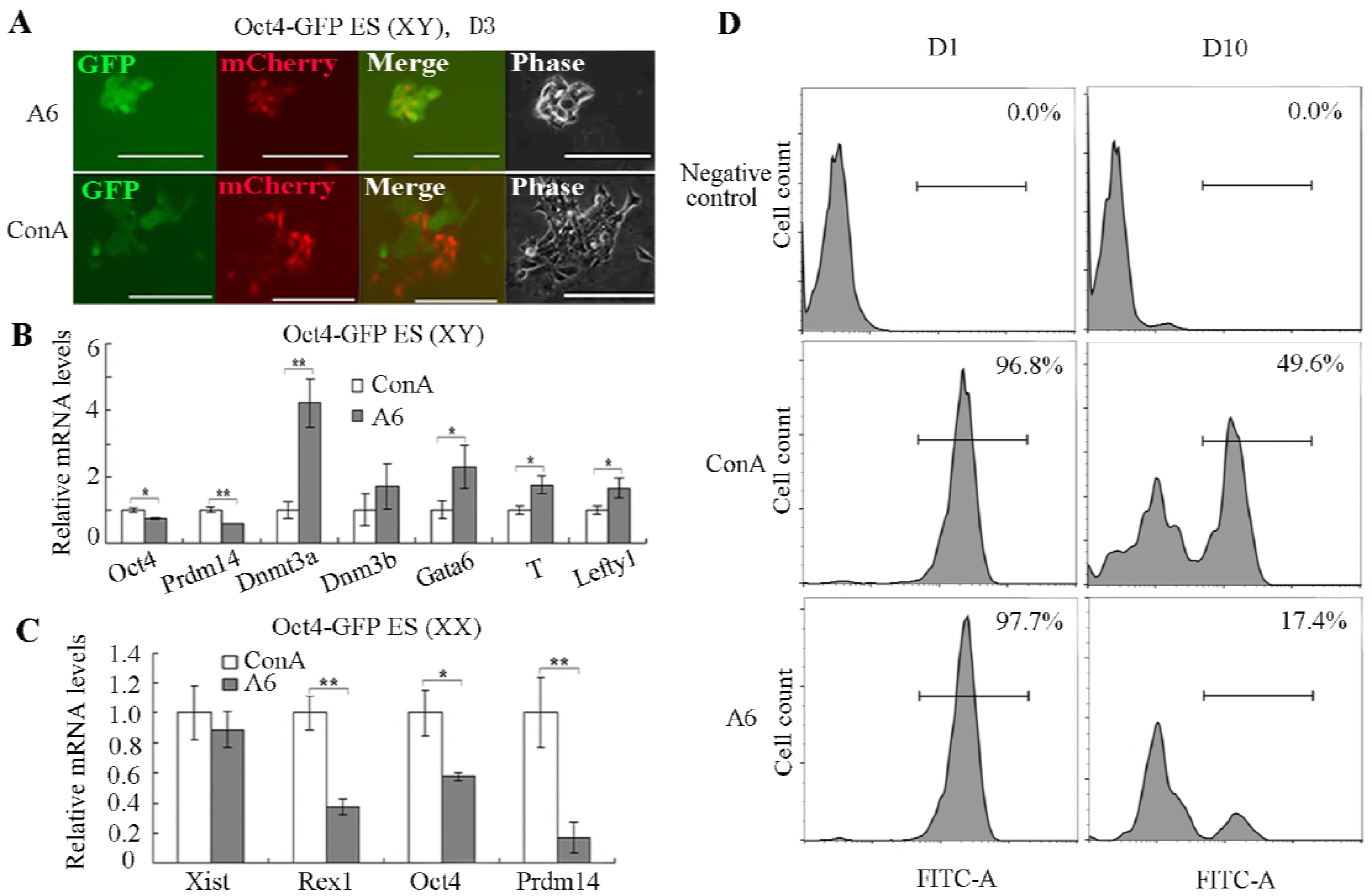
A6 accelerates ESC differentiation. (A) Images of Oct4-GFP ESCs expressing A6 or ConA for 3 days. Scale bars = 200 μm. (B and C) RNA levels of several genes in male and female ESCs expressing A6 on Day 5. (D) *Oct4*-GFP ESCs expressing A6 or ConA were analyzed for GFP expression in flow cytometry on days 1 and 10. Results are representative of three independent experiments and are mean ± SD. n = 3. **p <0.01. *p <0.05.

### The mechanism of R6 improving MEF reporgramming

Since the absence of *Xist* intron 1 enhancer has no obvious effects on iPSC formation from MEFs [22], the profound effects of R6 are unexpected and intriguing. One of the main co-repressors of KRAB is KRAB-Associated protein 1 (KAP-1), which can recruit the chromatin modifier SETDB1 for increasing H3K9me3 levels [35–39]. KAP1 also binds heterochromatin protein 1 (HP1), which interacts with H3K9me3 and stabilizes the repressive complex in chromatin [40]. Also, it is reported that TALE-dTF repressors can alter histone modifications at the targeted genomic regions [25]. We thus examined the effects of R6 on several histone modifications at the *Xist*intron 1 enhancer. We collected MEFs expressing R6 or ConR, and performed chromatin immunoprecipitation qPCR (ChIP-qPCR) by using three independent pairs of primers near R6 binding site to determine the levels of histone modifications at the *Xist* intron 1 enhancer region (Figure EV4A and Table EV2). Among all the histone modifications examined (H3K9me3, H3K27me3, H3K4me1, H3K4me3 and H3K27ac), R6 expression caused substantially higher H3K9me3 (Figure 5A and Figure EV4B), which may alter nuclear architecture [41–43]. To validate this, we performed 3D DNA-FISH [44] to explore the actual separation between defined chromosome loci under our experiment. We designed three FISH probes in X chromosome such that one probe for *Xist* and the other two probes targeting the each side of the X chromosome, named X^up^ and X^down^ (Figure 5B). We then verified that the distances between probes (Xist-X^up^, Xist-X^down^ and X^up^-X^down^) were longer in female MEFs expressing R6 than those carrying ConR (Figure 5C and Figure 5D; P<0.01), and the similar observation was found in male MEFs (Figure EV4C and EV4D; P<0.01), which indicated that both active and inactive X chromosomes were opened by R6 binding. Another possible approach that R6 affects iPSC generation and SCNT is by directly or indirectly regulating pluripotent genes. To investigate this possibility, we collected MEFs expressing R6 or ConR for five days and examined expression of several pluripotent genes. Consistent with the effects on both MEF reprogramming to iPSC and SCNT embryos reconstruction, R6 expression alone could slightly activated the expression of the endogenous *Oct4* at detectable levels (Figure 5E and Figure EV4E; P<0.01). To investigate the link between the three-dimensional change of X chromosome and the up-regulation of pluripotency-related genes in R6-expressing MEFs, the chromosome 17, where the critical pluripotent gene *Oct4* is located, were selected to examine the 3D structure by performing DNA-FISH. Another three DNA-FISH probes in chromosome 17 were designed to target *Oct4* locus site and its upstream (O^up^) and downstream (O^down^) sequences, respectively (Figure 5F). Similar with the observation in structural alteration of X chromosomes, the compact chromosome 17 were decondensed in MEFs expressing R6 (Figure 5G), since the distance between probes (Oct4-O^up^, Oct4-O^down^ and O^up^-O^down^) were significantly increased in 3D space within R6 expression (Figure 5H; P<0.01).

**Figure 5.**
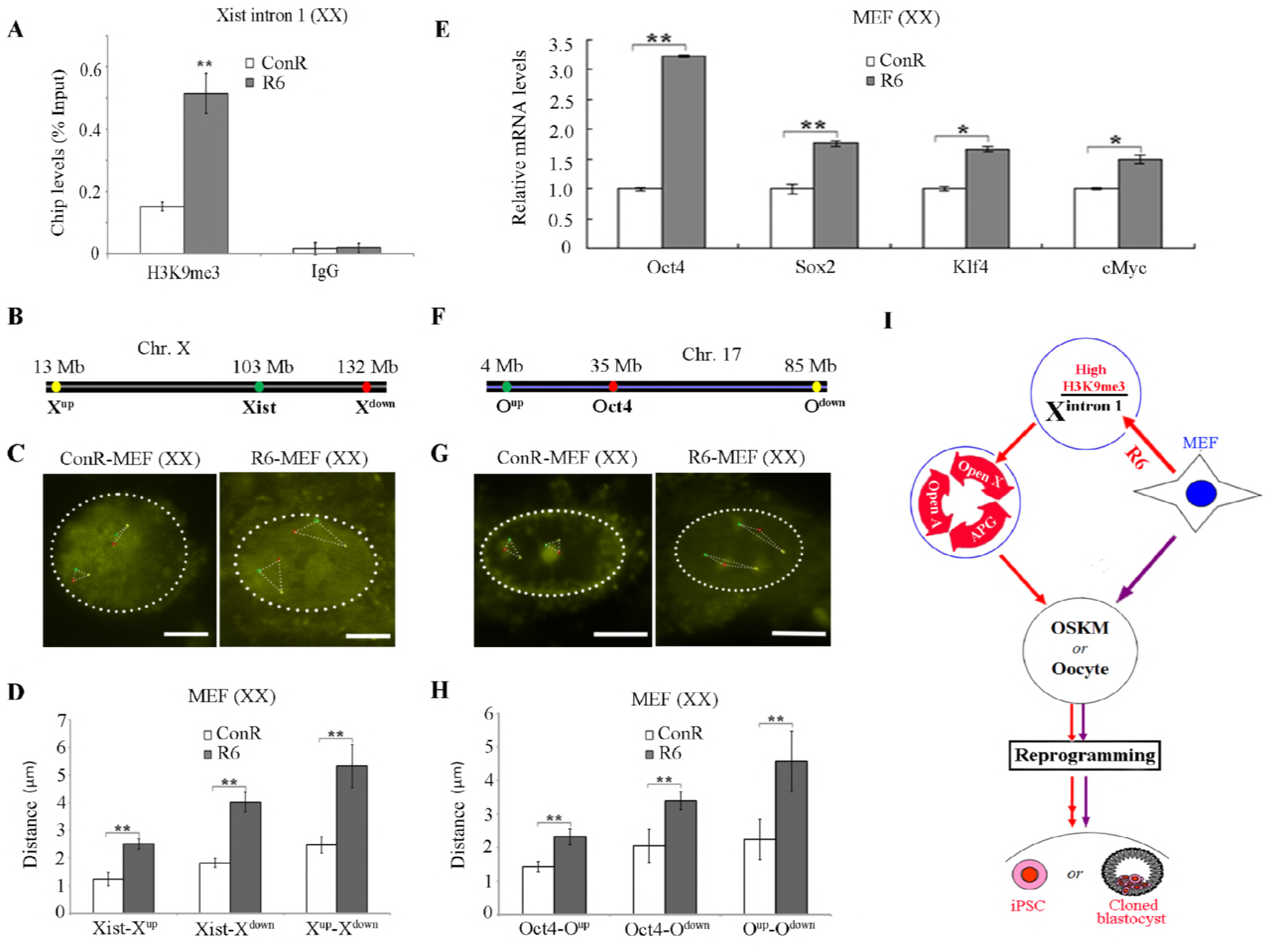
The mechanism of R6 improving MEF reporgramming. (A) ChIP-qPCR for H3K9me3, using primer pair 3 in female MEFs expressing ConR or R6. (B) Three DNA-FISH probes (X^up^, Xist, X^down^) were designed in X chromosome. (C) Representative DNA-FISH images of X chromosomes in female MEFs expressing R6 (right panel) and ConR (left panel), respectively. Scale bars=6μm. (D) The female MEFs expressing R6 had significantly different distances between probes (X^up^ - Xist, Xist - X^down^ and X^up^ - X^down^) from those carrying ConR. (E) qRT-PCR analysis of *Oct4, Sox2, Klf4* and *c-Myc* expression in female MEFs expressing R6 on Day 5. (F) Three DNA-FISH probes (O^up^, Oct4, O^down^) were designed in chromosome 17. (G) Representative DNA-FISH images of chromosome 17 in female MEFs expressing R6 (right panel) and ConR (left panel), respectively. Scale bars=6μm. (H) The different distances between probes (O^up^ – Oct4, Oct4 - O^down^ and O^up^ - O^down^) in female MEFs expressing R6 from those carrying ConR. (I) A diagram illustrating how R6 binding to *Xist* intron 1 affecting MEF reprogramming. Open X: Open X chromosomes; Open A: Open Autosomes, including chromosome 17; APG: Activation of Pluripotency-related Genes, such as *Oct4.* Results are representative of three independent experiments and are mean ± SD. n > 3. **p <0.01. *p <0.05.

In summary, our data support a model where R6 binding to the *Xist* intron 1 enhancer caused substantially higher H3K9me3, which opened X chromosomes, thus followed by autosomal structure altered in three dimensions, such as chromosome 17 (Figure 5I). Accompanying this dramatic chromatin decondensation were the alteration of chromatin-based activities, and the activation of pluripotent genes, such as *Oct4* (Figure 5I). Eventually, all those changes induced by R6 binding to the *Xist* intron 1 enhancer cooperatively improved both MEF reprogramming mediated by OSKM and oocytes (Figure 5I).

## Discussion

*Xist* regulates XCI in mouse female cells. The role of *Xist* in epigenome-resetting in somatic cell reprogramming to iPSCs has been suggested. In this study, we expressed and tested dTF repressors that are specific to several regions at the *Xist* locus. Surprisingly, the dTF repressor for the intron 1 enhancer region did not noticeably affect *Xist* expression levels in female MEFs but had profound positive effects on reprogramming MEFs to iPSCs. The dTF activator for the same genomic region substantially impeded the reprogramming. In contrast, the dTF for the promoter region of the *Xist* locus did not have such profound effects. These results are also in line with a previous study where knockout of the *Xist* intron 1 region did not affect *Xist* expression in MEFs or iPSC formation [22].

Dysregulation of gene expression, including *Xist*, is frequently found in SCNT embryos, which have low developmental efficiency and low cloning success rates. It has been reported that impeding *Xist* expression by either genomic deletion or RNAi knockdown increases birth rates of SCNT embryos, but the rate of SCNT preimplantation embryos is not substantially different [9, 10]. Remarkably, transient expression of R6 in both female and male MEFs substantially increased the developmental rates of SCNT morulae to blastocysts, which was consistent with the observation of ESC marker gene expression increased by R6. In the meantime, the results showed the effects of R6 on SCNT-embryo development are much more pronounced in male embryos than in females, which are in line with the previous reports about *Xist* effect on mouse SCNT that male reconstructed embryos are more sensitive and susceptible than females [10, 45]. Transcriptomic analysis of individual morula demonstrated that embryos reconstructed from R6-expressing MEFs clustered together with those morulae from IVF, and that R6 drastically reduced the commonly abnormally expressed X-linked and autosomal genes in SCNT embryos, compared to the control SCNT morulae. Also, Hierarchical clustering transcriptome analysis showed that the transcriptomes of male SCNT morulae carrying R6, which expressed lower level of *Xist* mRNA (FPKM=76), were more closed clustered with those of IVF embryos than female ones with higher *Xist* expression (FPKM=106; P<0.05), since R6 has no ability to down-regulate *Xist*, and abnormal expression of *Xist* prevented the cloned embryo development [9, 10]. This inference can be evidenced by the results that R4, which decreased the *Xist* expression in both femle MEFs and SCNT morulae, had the similar effects on transcriptomes of both female and male reconstructed morulae, compared with IVF embryos. However, R4 were much less effective than R6 in SCNT experiments, demonstrating that the effects of R6 on epigenome-resetting are largely independent of *Xist* transcript levels.

On the other hand, the dTF activator for the intron 1 enhancer blocked the iPSC generation and preimplantation development of SCNT embryos, which was in line with the ESC differentiation induced by A6. Those observations indirectly validate the role of R6 in improving MEF reprogramming.

R6 binding to the *Xist* intron 1 enhancer, which could be similar to in ESCs where the intron 1 enhancer is bound by multiple pluripotent and other factors, presumably recruiting repressive histone modifications [22], caused substantially higher H3K9me3. Consistent with previous reports [35–39], the enrichment of H3K9me3 in *Xist* intron 1 enhancer region remodeled the architecture of X chromosome to open state, and followed by the opening of closed autosomes, including chromosome 17.

Accompanying this dramatic chromatin decondensation was the alteration of chromatin-based activities, which led to the activation of pluripotent genes, such as *Oct*4. All those changes induced by R6 binding to the *Xist* intron 1 coordinately promoted both MEF reprogramming mediated by OSKM and oocytes.

In summary, our results have revealed an unexpected role of the *Xist* locus, in particular the intron 1 enhancer, in iPSC induction and SCNT embryo development. The new information provides a basis for improving reprogramming efficiency by further manipulating the *Xist* locus.

## Materials and Methods

### Mice

Housing and breeding of mice, and experimental procedures using mice were conducted in accordance with the UK 1986 Animals Scientific Procedure Act and the local institute ethics committee regulations.

### Plasmid vector construction

To create PB-TRE-*Oct4* and PB-TRE-CKS vectors, the TRE promoter was amplified from pTight (clontech) and cloned into a PB-bqA vector. cDNA of the mouse *Oct4* was cloned into PB-TRE transposon vectors, and the cDNAs of *c-Myc, Klf4* and *Sox2* were tandemly cloned into PB-TRE transposon vectors. *Xist* dTFs were reconstructed from *Oct4* dTF as reported in a previous study (Gao et al. 2013) by exchanging the binding domain of TALE protein (Table EV1).

### Preparation and transfection of MEFs

The MEFs were derived from E13.5 mouse embryos with a mixed 129S5/C57B6J background and cultured in M10 medium. Knockout DMEM (Invitrogen), 10% fetal bovine serum (Hyclone), 1 × glutamine-penicillin/streptomycin (Invitrogen) and 1 × nonessential amino acids (NEAA; Invitrogen) were included in this medium. MEFs were transfected by Amaxa Nucleofector (Lonza) using program A-023, and the transfection efficiency is about 8%.

### Flow cytometry

Mouse cells growing in 6-well plates were trypsinized and resuspended in M15 medium. The mixture was centrifuged at 200×g (Eppendorf centrifuge 5702R, A-4-38 rotor) for 3 min, and the medium was removed by inverting and plated onto tissue paper. Transgenic cells were resuspended in PBS and analyzed by Cytomics FC-500 (Bechman Coulter).

### Quantitative real-time PCR

RNA was isolated using the RNeasy Mini Kit (Qiagen). The samples were subsequently quantified and treated with gDNA wipeOut. First-strand cDNA was prepared by using the QuantiTect Reverse Transcription Kit (Qiagen). For each RT-PCR, 50 to 100 ng of cDNA were used for polymerase chain reaction (PCR) amplification. Standard PCR conditions were: 94 °C for 30s, 60 °C for 30s, and 68 °C for 30s for 30 cycles. For real-time PCR, we used TaqMan Gene Expression Assays. Taqman probes were purchased from Applied Biosciences (Table EV4). All quantitative PCR was performed in a 9700HT Fast Real-Time PCR System (Applied Biosciences). Mouse gene expression was determined relative to mouse GAPDH using the ΔCt method.

### ChIP analysis

10 million ESCs were cultured in a 10-cm, or 100 million MEFs were cultured in a 15-cm, and were collected 3 days after transfection of TALE-expressing plasmids and trypsinized for 5 min; trypsin was quenched by adding 10 ml media containing 10% FBS. The cell suspension was diluted to 40 ml with PBS and the cells were fixed for 12 min in formaldehyde at a final concentration of 1%. Cell crosslinking was quenched by adding 2.5 M glycine (0.125 M final concentration) before the cells were incubated on ice. Crosslinked cells were spun at 600 × g for 5 min and nuclei were prepared by consecutive washes with P1 buffer [10 mM Tris pH 8.0, 10 mM EDTA (pH 8.0), 0.5 mM EGTA, 0.25% Triton X-100] followed by P2 buffer (10 mM Tris pH 8.0, 1 mM, EDTA, 0.5 mM EGTA, 200 mM NaCl). Cell pellets were resuspended in 2 ml of ChIP lysis buffer [50 mM HEPES/KOH, pH=7.5, 300 mM NaCl, 1 mM EDTA, 1% Triton X-100, 0.1% DOC, 0.1% SDS, protease inhibitors complete mini (Roche)] and then sonicated using BioRuptor (Diagenode) in which they were pulsed for 15 cycles, each of 30 s sonication and 30 s rest. DNA was sheared to a size range of 500 - 1000 bp (confirmed on agarose gel). IgG (Cell Signalling, 2729S), antibodies for the HA tag (Abcam) and H3K9me3 (Abcam) were used in ChIP analysis. The primers for qRT-PCR are listed in Table EV2.

### iPSC induction and culture

To reprogram MEFs, vectors (in most experiments, 2.0 μg PB transposon, 2.0 μg 4F plus 2.0 μg *Xist* dTF, and 1.0μg PB transposase plasmid) were first mixed with 1 × 10^6^ cells in Opti-MEM (Invirogen), and the cells were electroporated with Amaxa Nucleofector (Lonza). Next, the cells were plated onto gelatinized 10 cm dishes in M10 for recovery over 24 h. The cells were then washed with phosphate buffered saline (PBS) and switched to M15 medium with Dox: knockout DMEM (Invitrogen), 15% FBS (HyClone), 1 × glutamine-penicillin/streptomycin (Invitrogen), 1 × nonessential amino acids (Invitrogen), 0.1 mM 2-mercaptoethanol (2-ME; Sigma), and 10^6^ U/ml LIF (Millipore). The medium was changed every other day, and the emerging iPSC colonies were monitored under a microscope. On day 14, the medium was changed to 2i/LIF medium [46, 47] with slight modifications [DMEM/F12 (Invitrogen), 1 × L-glutamine-penicillin/streptomycin (Invitrogen), N2, B27 (Invitrogen), 2-mecaptoethanol (2-ME; Sigma), PD (1.0 μM), CH (3.0 μM), and LIF (Millipore)], in which the cells were cultured further.

### Preparation of donor cells for SCNT

Mouse embryos were isolated from E13.5 C57BL/6J females mated to males carrying a mixed background of MF1, 129/sv and C57BL/6J strains. Both male and female MEFs were co-transfected by *Xist* dTF repressor and a plasmid carrying the Puro-IRES-GFP cassette under the control of the CAG promoter, then cultured for 2 d with Dox induction. Puromycin (2.0 μg/ml; Sigma) was added to the culture medium for a further 3 d to select for the transgenic cells. Two days after puromycin withdrawal, the surviving MEFs, both GFP and mCherry positive, were used as donor cells.

### Nuclear transfer and cloned embryo culture

Nuclear transfer was carried out as described [48, 49]. Briefly, recipient oocytes were collected from superovulated BDF1 female mice and enucleated in M2 medium (Sigma) containing 7.5 μg/ml cytochalasin B (Sigma). Thereafter, the donor cells were injected into the perivitelline space of the enucleated oocytes using a Piezo-driven micromanipulator (PiezoXpert; TransferMan NK2). The donor cells were electrofused with enucleated oocytes by cell fusion instrument (CF-150B; BLS). The reconstructed embryos were cultured for 2 h before the reconstructed embryos were activated in KSOM (Millipore) medium containing 2mM EGTA (Sigma), 5mM SrCl_2_ (Sigma) and 5μM Lata (Sigma) for 6 h, followed by further culture in M16 medium (Sigma) without added Dox. After 48 h, the cloned 2-cell embryos were cultured in KSOM for 96h, before being statistically assessed for pre-implantation development (Figure EV3A).

### Expression analysis of single morula samples

Reads pass quality control (over 50% bases with quality value >5 and less than 10% bases undetermined) were mapped to mm10 with Tophat (version 2.0.8). Then the gene expression level was calculated by cufflinks (version 2.2.1) and normalized to Fragments Per Kilobase of exon model per Million mapped reads (FPKM).

### Single morula RNA-seq library construction and sequencing

Single Morulae were picked into 5μ1 lysis buffer using micromanipulation. The samples were then amplified following the published protocol [50]. Hundreds of nanograms of cDNA were thus obtained after 20 cycles of PCR amplification.Subsequently, 300 ng cDNA was sonicated with Covaris S220 and then used for library construction employing the NEBNext^®^ Ultra™ DNA Library Prep Kit for Illumina. Libraries were then sequenced on an Illumina Hiseq2500 platform for 100 bp pair-end reads.

### Clustering and PCA analysis

The top 200 genes that showed differential expression among all the samples were picked out for further principle component analysis and clustering using ggplot2 and heatmap2 in R studio (version 3.0.1).

### 3D DNA-FISH

Three DNA-FISH probes were designed in X chromosome (X^up^: 13280496-13293988 bp; Xist: 103460373-103483233 bp; X^down^:136203771-136225714 bp) and chromosome 17 (O^up^: 4000853-4009896 bp; Oct4: 35501777-35510777 bp; O^down^: 85000474-85009474 bp). The target DNA products labeled with haptens-biotin or digoxigeni were amplified by PCR. Labeled PCR products were dissolved in a standard hybridization buffer (50 % deionized formamide (ICN), 2× saline-sodium citrate (1× SSC: 0.15 M NaCl, 15 mM Na_3_C_6_H_5_O_7_), 10 % dextran sulphate (Pharmacia Biotech) to a final concentration of 20–40 ng/μ1 with a 50-fold excess of DNA. We collected MEFs expressing ConR or R6 by flow cytometry, and performed multicolor FISH on 3D-preserved nuclei as described [44]. To ensure full probe penetration into cell nuclei, the optional pepsin treatment was included. Nuclei were counterstained with 4’, 6-diamidino-2-phenylindole. Images were acquired using an LSM710 confocal microscope (Zeiss). Images were analyzed and distances were measured using the LSM Image Browser (Carl Zeiss) and Imaris software (Bitplane).

### Statistical analysis

The data are presented as the mean ± standard deviation (SD). Significance was determined using the Student’s unpaired ttest with two-tailed distribution. p-Values < 0.05 were considered significant.

## Acknowledgements

We thank Beiyuan Fu and Fengtang Yang for their assistance with *Xist* RNA FISH, and are grateful to Jitong Guo for the micromanipulation expertise. The work was supported by the Ministry of Science and Technology project of Inner Mongolia (No. 20130216), the National Natural Science Foundation of China (No. 31560335), and by the Wellcome Trust (grant number 098051).

## Author contributions

J.Z. designed and did most of the experiments, analyzed and interpreted data, made all figures, and contributed to the writing of the manuscript; X.G., J.Y., X.F., J.T., W.W., Y.L., L.F., H.H., X.X., F.T. and S.B. did experiments, provided reagents, or provided intellectual input; and P.L. and X.L. designed the experiments, supervised the research, and wrote the manuscript.

## Expanded View Figure legends

**Figure Ev1**. Assembly and functional assessment of *Xist* dTFs. Related to Figure 1. (A) A schematic diagram for TALE dTFs in the PB delivery vector. (B, C) FISH with DNA probes targeting *Xist* RNA (green) in female R4-transfected MEFs and WT MEFs on D5. Scale bars=10µm.

**Figure EV2.** iPSCs produced with various combinations of *Xist* dTF and OSKM. Related to Figure 2.

(A) Schematic for the generation of iPSCs by 4F +/− Xist dTF. (B) In vitro differentiation of established iPSCs. Scale bars=100µm. (C) Reprogrammed iPSC colonies were visualized by AP staining. (D, E) Quantitation of female and male *Rex1*-GFP iPSC colony induced by 4F+dTFs. Results are representative of three independent experiments and are mean ± SD. n = 3. **p <0.01.

**Figure EV3**. Xist dTF repressors improve cloned blastocyst development *in vitro*. Related to Figure 3.

(A) Schematic for the reconstructed embryo generation. (B, C) The percentage of female and male cloned embryos that reached the indicated stages. (D, E) PCA of amplified RNAs from single male and female cloned morulae, and *in vitro* fertilization (IVF)-derived embryos were used as positive control. Results are representative of three independent lines and are mean ± SD. n = 3. **p <0.01.

**Figure EV4.** The mechanism of R6 improving MEF reporgramming. Related to Figure 5.

(A) Schematic for the ChIP-qPCR primer position at the *Xist* intron 1. (B) ChIP-qPCR for H3K9me3 in male MEFs expressing ConR or R6. (C) Representative DNA-FISH images of X chromosome in male MEFs expressing R6 (right panel) and ConR (left panel), respectively. Scale bars=6µm. (D) The different distances between probes (Xist-Xup, Xist-Xdown and Xup-Xdown) in male MEFs expressing R6 from those carrying ConR. (E) qRT-PCR analysis of *Oct4, Sox2, Klf4* and *c-Myc* expression in male MEFs expressing R6 on Day 5. Results are representative of three independent experiments and are mean ± SD. n ≥3. **p <0.01.

